# Design of a novel multi-epitope mRNA vaccine against BtHKU5-CoV-2 using immunoinformatics

**DOI:** 10.1101/2025.09.02.673669

**Authors:** Ningze Zheng, Yingqi Xu

## Abstract

Bat HKU5-CoV-2 (BtHKU5-CoV-2), a recently discovered bat-infecting merbecovirus, was found to infect human cell lines by utilizing the human angiotensin-converting enzyme 2 (ACE2) receptor, similar to SARS-CoV-2, which caused millions of deaths. Moreover, its broad host tropism has raised significant concerns about potential human spillover risk. Therefore, there is an urgent need to develop vaccines to combat the potential outbreak of BtHKU5-CoV-2. However, research focusing on BtHKU5-CoV-2 remains limited. In this study, we designed a novel multi-epitope vaccine against BtHKU5-CoV-2 using an immunoinformatic approach. Eight cytotoxic T lymphocyte (CTL) epitopes, seven helper T lymphocyte (HTL) epitopes, and five linear B lymphocyte (LBL) epitopes were screened from the spike glycoprotein of BtHKU5-CoV-2. The selected epitopes were joined together with an appropriate linker, and β-defensin II and MHC I-targeting domain (MITD) were incorporated into the construct to enhance vaccine immunogenicity. Biological characteristic analysis revealed that the designed vaccine exhibited strong antigenicity and immunogenicity while being non-toxic and non-allergenic. The tertiary structure of the multi-epitope vaccine was modeled, refined, and validated, demonstrating its structural stability and near-native conformation. Molecular docking studies showed that the vaccine successfully docked with Toll-like receptor 2 (TLR2) and TLR4. Moreover, its mRNA exhibits strong interactions with TLR3, TLR7, and TLR8 receptors. Additionally, *in silico* immune simulations have suggested that vaccination could trigger robust humoral and cellular immunity. These findings suggest that the proposed mRNA vaccine is a potential candidate for targeting BtHKU5-CoV-2. Further experiments are necessary to validate its protective efficacy.

**Author summary:** BtHKU5-CoV-2, a newly discovered merbecovirus isolated from bats, exhibits potential for spillover into humans. It was found to utilize human ACE2 as functional receptors for infection. A functional receptor acts like a “key” that fits into the “lock” on the host cell, enabling viral entry. BtHKU5-CoV-2 warrants significant attention, because it shares the same functional receptor with SARS-CoV-1 and SARS-CoV-2, which caused the 2003 SARS epidemic and the 2019 pandemic, respectively. Thus, developing vaccines to prevent potential global outbreaks of BtHKU5-CoV-2 is urgently needed. Theoretically, within the body’s immune surveillance system, proteins from BtHKU5-CoV-2 are processed via proteasomal degradation into short peptides. The peptides with immunogenicity bind to MHC molecules and are presented on the cell surface. These peptides, known as epitopes, can initiate immune reaction. In this study, we designed a multi-epitope mRNA vaccine against BtHKU5-CoV-2 using immunoinformatics methods. Epitopes were screened from the spike glycoprotein, a promising target of BtHKU5-CoV-2. Our results suggest that the vaccine is safe and capable of inducing strong humoral and cellular immunity. Therefore, this mRNA vaccine represents a promising candidate for preventing furture BtHKU5-CoV-2 outbreak.

## Introduction

BtHKU5-CoV-2, a positive-strand RNA virus, was first reported in *Pipistrellus* bat anal swab samples collected from several provinces of China in 2025 [1]. It is a new merbecovirus belonging to the same family as other coronaviruses that infect humans, such as MERS-CoV. Like other coronaviruses, BtHKU5-CoV-2 employs spike glycoprotein to recognize receptors and achieve the fusion of viral and cellular membranes [2]. More importantly, it was found to infect human cell lines by utilizing angiotensin-converting enzyme 2 (ACE2) as a functional receptor [1]. ACE2 is a known receptor of SARS-CoV-2, a zoonotic virus that has led to COVID-19 and caused millions of deaths in recent years [3, 4]. Additionally, BtHKU5-CoV-2 may infect various mammalian and avian species via ACE2 orthologs [1]. Therefore, BtHKU5-CoV-2 has the potential for zoonotic transmission and threatens public health, raising immediate concerns [5–7].

mRNA vaccines play crucial roles in reducing hospitalizations and deaths caused by SARS-CoV-2 during the COVID-19 pandemic [8]. Thereafter, mRNA vaccine technology has rapidly developed and has become a frontline measure for defense against infectious diseases [9]. Compared to traditional recombinant protein vaccines and DNA vaccines, mRNA vaccines exhibit multiple advantages, including: (i) they are hardly incorporated into the host’s genome; (ii) mRNA can be produced in an in vitro cell-free system without complex purification procedures; (iii) multiple antigens can be inserted into one mRNA sequence; (iv) Unlike DNA vaccines, mRNA vaccines do not induce an immune response against components of the vector [10]. Overall, mRNA vaccines are fast, cheap, and flexible tools to defend against rapid outbreaks of infectious diseases.

Advancements in immunoinformatic approaches and the availability of immunological datasets have helped researchers screen target epitopes and design epitope-based vaccines more efficiently. In particular, multiple-epitope vaccines have gained great attention for rapidly mutating pathogens. On the one hand, researchers can identify potent conserved epitopes from viral components in a short time. On the other hand, cytotoxic T lymphocyte (CTL) epitopes, helper T lymphocyte (HTL) epitopes, and B cell epitopes can be incorporated into a multiple-epitope vaccine, resulting in simultaneous activation of cellular and humoral immune responses, which are necessary for defense against infectious pathogens [11]. Moreover, inclusion of adjuvants and components that assist in epitopes presentation boosts immunogenicity and extends immune responses [12, 13]. In recent years, some studies have successfully designed multi-epitope vaccines targeting different viruses and achieved the desired protective effects, such as mRNA vaccines against monkeypox virus [14] and HIV mRNA vaccines [15].

Some researchers have become aware of BtHKU5-CoV-2 because of its spillover potential. Therefore, it is necessary to develop an effective vaccine to combat rapid viral outbreaks. Combining mRNA platforms and epitope design is an ideal strategy for developing a multi-epitope mRNA vaccine against BtHKU5-CoV-2. In this study, we predicted the conserved CTL, HTL, and B cell epitopes from the spike glycoprotein of BtHKU5-CoV-2. We selected promising epitopes for the multi-epitope vaccine development. Immunogenicity, tertiary structure, interaction with immune receptors, and physicochemical properties of the vaccine were estimated in our study. Overall, the multi-epitope mRNA vaccine designed in the present study is a promising prophylactic and therapeutic agent to address potential pandemics of BtHKU5-CoV-2.

## Methods

### 1. Retrieval of target protein sequences

The amino acid sequences of the spike glycoproteins of BtHKU5-CoV-2-441 (C_AAI84074.1), BtHKU5CoV-2-153 (C_AAI84049.1), BtHKU5-CoV-2-155 (C_AAI84054.1), BtHKU5-CoV-2-023 (C_AAI84059.1), BtHKU5-CoV-2-028 (C_AAI84064.1), and BtHKU5-CoV-2-381 (C_AAI84069.1) were obtained from GenBase. Among these bat HKU5-CoVs, they share 91.2%-99.2% similarity in amino acid sequences, and BtHKU5-CoV-2-441 showed the highest identity with the others [1]. We chose BtHKU5-CoV-2-441 (referred to as BtHKU5-CoV-2 in the following analysis) as the representative strain for further study.

### 2. Epitopes prediction

#### 2.1. CTL epitopes prediction

To screen out the CTL epitopes of the target protein, NetCTL 1.2 (https://services.healthtech.dtu.dk/services/NetCTL-1.2/) was employed. In this step, a threshold of 0.75 (default value) was applied, while C-terminal cleavage, TAP transport efficiency, and peptide-MHCI binding were also considered. Twelve supertypes, A1, A2, A3, A24, A26, B7, B8, B27, B39, B44, B58, and B62, were considered in the screening process [16]. Antigenicity prediction of epitopes was subsequently performed using the Vaxijen 2.0 tool (https://ddg-pharmfac.net/vaxijen/VaxiJen/VaxiJen.html) with a threshold of 0.4 [17]. To ensure the selected epitopes were non-toxic and non-allergenic, AllerTOP 2.0 (http://www.ddg-pharmfac.net/AllerTOP) and ToxinPred 2.0 (https://webs.iiitd.edu.in/raghava/toxinpred2/) were exploited to exclude the epitopes with toxicity or allergenicity [18, 19]. The immunogenicity of epitopes is critical for activating the immune response; therefore, the epitopes were submitted to the Class I Immunogenicity server in the IEDB platform (http://tools.iedb.org/immunogenicity/) [20]. Immunogenic epitopes (values exceeding 0) were included in this study. Finally, to screen out which MHC I alleles that epitopes were capable of binding to, the TepiTool in the IEDB platform (http://tools.iedb.org/tepitool/) was applied, and the IC50 of selected epitopes should be less than 500 nM [21].

#### 2.2. HTL epitopes prediction

Prediction of HTL epitopes was conducted using the NetMHCIIpan - 4.0 server (https://services.healthtech.dtu.dk/services/NetMHCIIpan-4.0/), which uses Artificial Neural Networks (ANNs) to predict peptides that are capable of binding to indicated MHC II molecules [22]. In this screening process, HLA-DRB1 alleles (including 0101, 0301, 0401, 0404, 0701, 0802, 0901, 1101, 1302, 1501), HLA-DRB3_0101, DRB4_0101, DRB5_0101, HLA-DQA10501 (-DQB10201, -DQB10301), HLA-DQA10301-DQB10302, HLA-DQA10401-DQB10402, HLA-DQA10101-DQB10501, HLA-DQA10102-DQB10602, HLA-DPA10201-DPB10101, HLA-DPA10103 (-DPB10201, -DPB10401), HLA-DPA10301-DPB10402 and HLA-DPA10201 (-DPB10501, -DPB11401) were selected for prediction. Only epitopes with an IC50 < 500 nM were subjected to further analysis. The antigenicity, allergenicity, and toxicity of epitopes were characterized using vaxijen2.0, Allertop 2.0, and ToxinPred 2.0, respectively. Epitopes with an antigenicity score over the threshold of 0.4, while are non-toxic and non-allergenic will be considered. Finally, the epitopes’ potential ability to induce IL-2, IL-4, and IFN-γ was evaluated using IL2Pred (https://webs.iiitd.edu.in/raghava/il2pred/), IL4Pred (https://webs.iiitd.edu.in/raghava/il4pred/) and IFNepitope (https://webs.iiitd.edu.in/raghava/ifnepitope/) [23, 24]. The epitopes with IL-2-, IL-4-, and IFN-γ-inducing capacity were included in the following study.

#### 2.3. Linear B cell epitopes prediction

For predicting linear B cell epitopes (LBL), we employed the ABCpred tool (https://webs.iiitd.edu.in/raghava/abcpred/) in this study [25]. In this step, the default settings, a threshold of 0.51 and 16 amino acids as epitope length were used for screening. Subsequently, antigenicity, allergenicity, and toxicity of the epitopes were assessed using Vaxijen2.0, Allertop 2.0, and ToxinPred 2.0, respectively. Epitopes that were predicted to have the characteristics of an antigenicity score higher than 0.4, non-toxic, and non-allergenic, were chosen for vaccine construction.

### 3. Epitope conservation analysis and homology study

The complete genomes of all six bat HKU5-CoV lineage 2 (HKU5-CoV-2) strains were downloaded from GenBase following accession numbers: C_AA085189.1, C_AA085190.1, C_AA085191.1, C_AA085192.1, C_AA085193.1, and C_AA085194.1. The nucleotide sequence of spike glycoprotein was extracted from the corresponding genomic sequence. Conservation analysis of epitopes across the six bat HKU5-CoV-2 strains was conducted using the local *tblastn* tool. The epitopes that were highly conserved or lacking mutation sites were considered in the following study. To avoid cross-immunity or tolerance to epitopes, the complete human proteome and complete proteome of seventy-nine bacterial species that commonly reside in the human gut were downloaded from the NCBI database. A local database was established using Diamond software, and then the BLASTp program was employed to analyze the homologs between the candidate epitopes and human proteomes, as well as intestinal bacterial proteomes. The selected epitopes met the following criteria: E-value exceeds 10^-4^ and a bit score less than 100.

### 4. Multi-epitope vaccine construction and physicochemical properties prediction

Following screening, epitopes with strong antigenicity, non-toxicity, and non-allergenicity were introduced in vaccine design. For joining the epitopes, AAY linkers were used for CTL epitopes, GPGPG linkers were used for HTL epitopes, while KK linkers were used for LBL epitopes. In addition, β-defensin II, an immunological adjuvant, was attached to the N-terminus of the sequences using the EAAAK linker. A TAT peptide, which is capable of assisting the vaccine traversing the cell membrane, was added to the C-terminal of the vaccine and connected by an EAAAK linker. Finally, the ProtParam tool (https://web.expasy.org/protparam/), Vaixijen 2.0, ToxinPred2.0 and AllerTOP 2.0 were employed to predict the physicochemical properties, antigenicity, toxicity, and allergenicity of the vaccine, respectively.

### 5. Structure modeling of the vaccine

The secondary structure of vaccines was predicted using PISPRED 4.0 (https://bioinf.cs.ucl.ac.uk/psipred/), and elements such as strands, helices, and coils are shown in the structure. The tertiary structure of vaccines was modeled using the Robetta service (https://robetta.bakerlab.org/), which predicts the protein structure based on an accurate deep learning-based method, RoseTTAFold [26]. Subsequently, the GalaxyRefine server (https://galaxy.seoklab.org/cgi-bin/submit.cgi?type=COMPLEX) was used for structural optimization of the models [27]. Only the refined model with lower MolProbity scores underwent quality estimation using the ProSA-Web (https://prosa.services.came.sbg.ac.at/prosa.php), PROCHECK in the PDBsum platform, and ERRAT [28]. Visualization of the 3D structure of vaccines was performed using PyMOL v2.5 software.

### 6. Conformational B cell epitopes prediction

Spike glycoprotein-specific antibodies play a significant role in the prevention of coronavirus infection. The possible conformational B-cell epitopes contained in vaccine structures were predicted using ElliPro (http://tools.iedb.org/ellipro/), an antibody epitope prediction server [29]. The tool predicts discontinuous antibody epitopes according to the 3D shape of the protein and outputs a score (protrusion index, PI) for each predicted epitope.

### 7. Molecular docking

Two molecules were favored for complex docking: TLR2 and TLR4, which are important pattern recognition receptors in mammals. TLR2 and TLR4 can trigger innate antiviral responses via direct or indirect interactions with viral glycoproteins [30]. From the RCSB PDB database, the crystal structures of TLR2 and TLR4 were downloaded with PDB ID of 2Z7X and 4G8A, respectively. After cleaning, such as removing non-protein components, the tertiary structures were submitted to the ClusPro 2.0 server (https://cluspro.org/home.php) for molecular docking [31]. Subsequently, the structures of the complexes were refined using the HADDOCK 2.4 server (https://rascar.science.uu.nl/haddock2.4/), which drives the docking process based on information from predicted protein interfaces in ambiguous interaction restraints (AIRs). Finally, the detailed interaction between the components in the complexes was illustrated using the PDBsum server (https://www.ebi.ac.uk/thornton-srv/databases/pdbsum/Generate.html).

### 8. Molecular dynamic simulation

Molecular dynamics (MD) simulations were conducted using the iMODs tool (https://imods.iqf.csic.es/) to evaluate the stability of the docking complex [32]. The online server explores the collective motion of proteins in internal coordinates using the normal modal analysis (NMA) method. Indicators, including flexibility and covariance, were used to assess the stability of vaccine/TLR complexes.

### 9. Population coverage calculation and immune simulation analysis

Only in individuals who express specific HLA molecules that bind to and present corresponding epitopes on the cell surface can the epitope trigger an immune response. The expression and distribution of HLA subtypes varies in different counties and regions. Therefore, the Population Coverage server in the IEDB platform (http://tools.iedb.org/population/) was implemented to calculate the population coverage across the world for epitope sets contained in the vaccine. In this program, the calculation options, class I and class II combined, were selected for analysis in the present study.

To predict host’s immune response against vaccination, the C-IMMSIM webserver (https://kraken.iac.rm.cnr.it/C-IMMSIM/index.php?page=0) was employed to simulate the immune response profile following three vaccine injections [33]. HLA alleles, including HLA-A0201, HLA-A0231, HLA-B5301, HLA-B1501, HLA-DRB1_0405, and HLA-DRB1_0701, were selected for analysis. The duration of the simulation was 1000-time steps, which is approximately 350 days.

### 10. Construction of mRNA vaccine

In the final construction of the multi-epitope vaccine, the signal peptide of Tissue Plasminogen Activator, which enables the synthesized peptides to be secreted into the extracellular space, was added to the N-terminus of the sequence. Furthermore, the sequence of the MHC I-targeting domain (MITD) was attached to the C-terminus to improve the epitope presentation. The amino acid sequence of the vaccine was submitted to the GenSmart Codon Optimization tool of GenScript for reverse translation and codon adaptation in Homo sapiens. Additionally, a Kozak sequence was added to the 5’ end, and a TAA codon was used to stop translation. To improve the stability and translation efficiency of the mRNA, a 5’-UTR (derived from cytomegalovirus immediate-early gene), 3’-UTR (derived from human growth hormone), and 120-nt polyadenylate tail were attached to the mRNA structure. Full-length DNA sequence was cloned into the pET-28a (+) vector using the GenSmart Design server on the GenScript platform. This vector was presumed to be linearized by BamHI restriction enzyme and used as a template for in vitro mRNA transcription.

### 11. mRNA-TLR molecules docking analysis

To assess the interaction between the mRNA of the vaccine and TLR molecules, the transcription tool was first utilized to transcribe the full DNA sequence, covering the region from the 5’-UTR to the poly(A) tail, into mRNA, and then the RNAfold tool (http://rna.tbi.univie.ac.at/cgi-bin/RNAWebSuite/RNAfold.cgi) was used to predict the secondary structure of the vaccine’s mRNA.

Endosomal TLRs, including TLR3, TLR7, and TLR8, specialize in recognizing the nucleic acids of pathogens and subsequently activate the host’s innate immune system [34]. Therefore, interactions between TLRs and the mRNA vaccine sequence were examined in the present study. The amino acid sequences of human TLR3, TLR7, and TLR8 were obtained from the UniProt database with the accession numbers O15455, Q9NYK1, and Q9NR97, respectively. RNA-protein interactions were predicted using the RPISeq tool (http://pridb.gdcb.iastate.edu/RPISeq/batch-rna.html), which outputs the results in the RF and SVM classifier forms. Finally, the crystal structure of TLR molecules that can bind to the mRNA vaccine was retrieved from the RCSB PDB database, and the modeling docking between TLR molecules and vaccine mRNA was performed using the HDOCK server.

## Results

### 1. Prediction of epitopes

Using several credible epitope prediction tools, we identified eight CTL epitopes (Table 1), seven HTL epitopes (Table 2), and five LBL epitopes (Table 3) in the spike glycoprotein sequence of the BtHKU5-CoV-2-441 strain. All epitopes showed high antigenicity, but were non-toxic and non-allergenic. Further analysis revealed that HTL epitopes could potentially induce IFN-γ, IL-2, and IL-4 production. Among the LBL epitopes with a high predicted score of antigenicity, only those that lie within the receptor-binding domain of spike glycoprotein were selected [2].

**Table 1.** CTL epitope screening results.

| Epitope | Start position | HLA molecule | Rank% | IC50(nM) | Antigenicity | Conservation |
| --- | --- | --- | --- | --- | --- | --- |
| KTYSNITIA | 62 | HLA-A*30:01 | 0.089 | 20.42 | 1.0509 | 6/6 |
| TYSNITIAM | 63 | HLA-A*24:02 | 0.55 | 411.35 | 1.2420 | 6/6 |
|  |  | HLA-A*23:01 | 0.857 | 453.21 |  |  |
| LPYFHNINY | 279 | HLA-B*35:01 | 0.015 | 5.3 | 1.0163 | 6/6 |
|  |  | HLA-B*53:01 | 0.112 | 96.24 |  |  |
| VQMTFVISV | 547 | HLA-A*02:06 | 0.009 | 2.59 | 1.1614 | 6/6 |
|  |  | HLA-A*02:01 | 0.034 | 5.03 |  |  |
|  |  | HLA-A*02:03 | 0.141 | 7.64 |  |  |
| VSIPTNFSF | 757 | HLA-B*58:01 | 0.021 | 5.93 | 1.1587 | 6/6 |
|  |  | HLA-B*57:01 | 0.069 | 24.48 |  |  |
|  |  | HLA-B*15:01 | 0.337 | 60.24 |  |  |
|  |  | HLA-A*32:01 | 0.124 | 83.44 |  |  |
|  |  | HLA-A*23:01 | 0.345 | 110.43 |  |  |
|  |  | HLA-B*35:01 | 0.359 | 159.06 |  |  |
|  |  | HLA-A*24:02 | 0.482 | 329.45 |  |  |
| IPTNFSFGL | 759 | HLA-B*53:01 | 0.126 | 112.83 | 1.8318 | 6/6 |
| MEAAYTASL | 925 | HLA-B*40:01 | 0.008 | 5.69 | 0.6652 | 6/6 |
| LASFAAIPF | 946 | HLA-B*35:01 | 0.029 | 8.76 | 0.8943 | 6/6 |

**Table 2.**
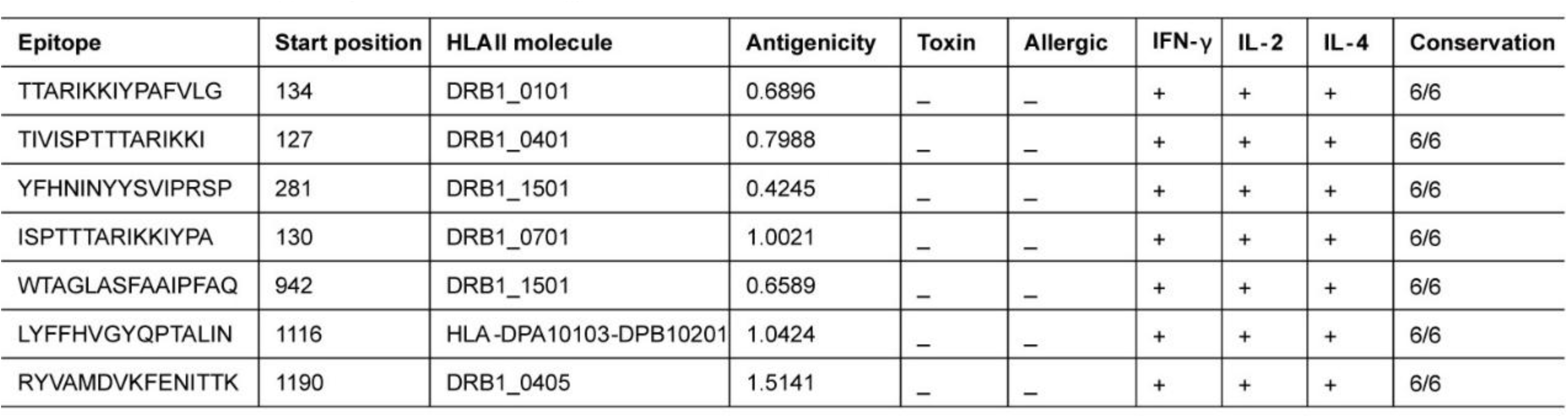
HTL epitope screening results.

**Table 3.**
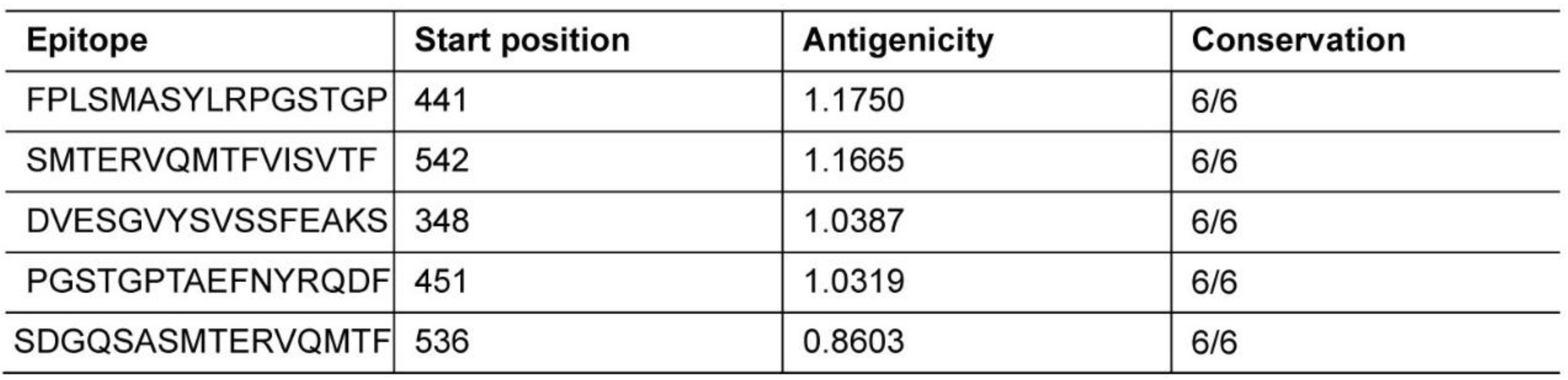
LBL epitope screening results.

By the mean of local *tblastn* tool, we analyzed the conservation of selected epitopes across the six BtHKU5-CoV-2 strains, and the results revealed that all epitopes showed evident conservation or complete identity. Therefore, the findings indicate that the vaccine designed against the BtHKU5-CoV-2-441 strain might also be effectively protected against the other five strains.

### 2. Vaccine construction and physicochemical properties assessment

In the development of vaccines, CTL epitopes, HTL epitopes, and LBL epitopes are joined by AAY, GPGPG, and KK linkers, respectively. Furthermore, β-defensin II was attached to the N-terminus, whereas a TAT peptide was added to the C-terminus (Fig. 1A). The vaccine is comprised of 419 amino acids and has a molecular weight of 44.37 kDa. The antigenicity of the vaccine was predicted by Vaixijen 2.0, and ANTIGENpro servers, and the scores were 0.7537 and 0.943474, indicating that the vaccine has considerable antigenic potential. Meanwhile, the analysis by ToxinPred2.0 and AllerTOP 2.0, implied that the vaccine showed no allergenicity and no toxicity. Using the ProtParam program, it was predicted that the vaccine’s theoretical isoelectric point value was 9.56, its instability index was 25.81, and its aliphatic index was 70.31, suggesting that the vaccine protein has the potential to be stable. The half-life of the vaccine was estimated to be 30 h in mammalian reticulocytes (in vitro), >20 h in yeast, and >10 h in *Escherichia coli*. In addition, the solubility value was calculated to be 0.515 using the Protein-Sol web tool, demonstrating that the vaccine has high solubility (S1 Fig). Overall, the vaccine showed potentially favorable biophysical and biochemical attributes, indicating that it deserves further development.

**Fig. 1.**
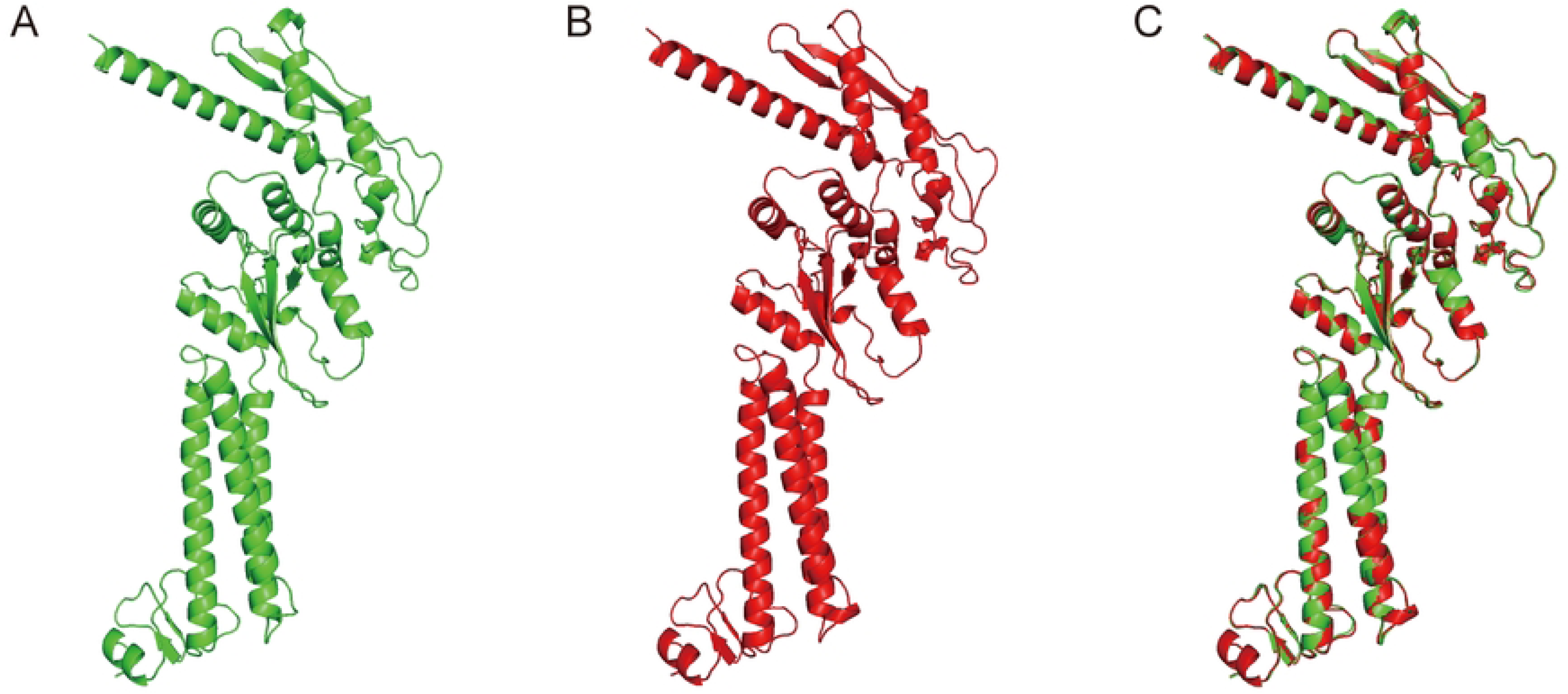
Schematic illustration of BtHKU5-CoV-2 vaccine construction. (A) The amino acid sequences of the vaccine. Each part is indicated by a different color, while β-defensin II adjuvant, CTL epitope, HTL epitope, LBL epitope, pan-HTL epitope and TAT peptide are labeled in green, blue, red, yellow, purple and orange, respectively. (B) The predicted secondary structure of the vaccine. The structure characteristics annotated with different color are illuminated by annotations below.

### 3. Prediction of secondary and tertiary structure

The secondary structure of the vaccine comprised 25.5% extended strand, 31% alpha helix, and 43.4% random coil, as predicted by the PISPRED server (Fig. 1B). The vaccine’s tertiary structure was modeled by the Robetta web tool, which predicts the 3D model of protein through Continuous Automated Model EvaluatiOn (CAMEO) (Fig. 2A). Subsequently, the three-dimensional structure of the vaccine was submitted to GalaxyRefine to refine the side chains, and a refined model with a Molprobity score of 2.006 is shown (Fig. 2B). An overlaid model of the original and refined constructs was generated and analyzed using PyMOL software (Fig. 2C). The quality of the refined construct was estimated using a Ramachandran plot, and the results showed that 90.1% of residues were in the favored region, 8.1% in the allowed region, and 1.7% in the disallowed region (Fig. 3A-B). Moreover, a Z-score of -7.54 was predicted by the Pro-SA web (Fig. 3C). The energy map showed that the energy values of most residues were less than zero (Fig. 3D). An overall quality score of 93.052 was generated using ERRAT (Fig. 3E). These results suggested that the refined tertiary structure of the vaccine obtained in this study exhibited good reliability and stability.

**Fig. 2.**
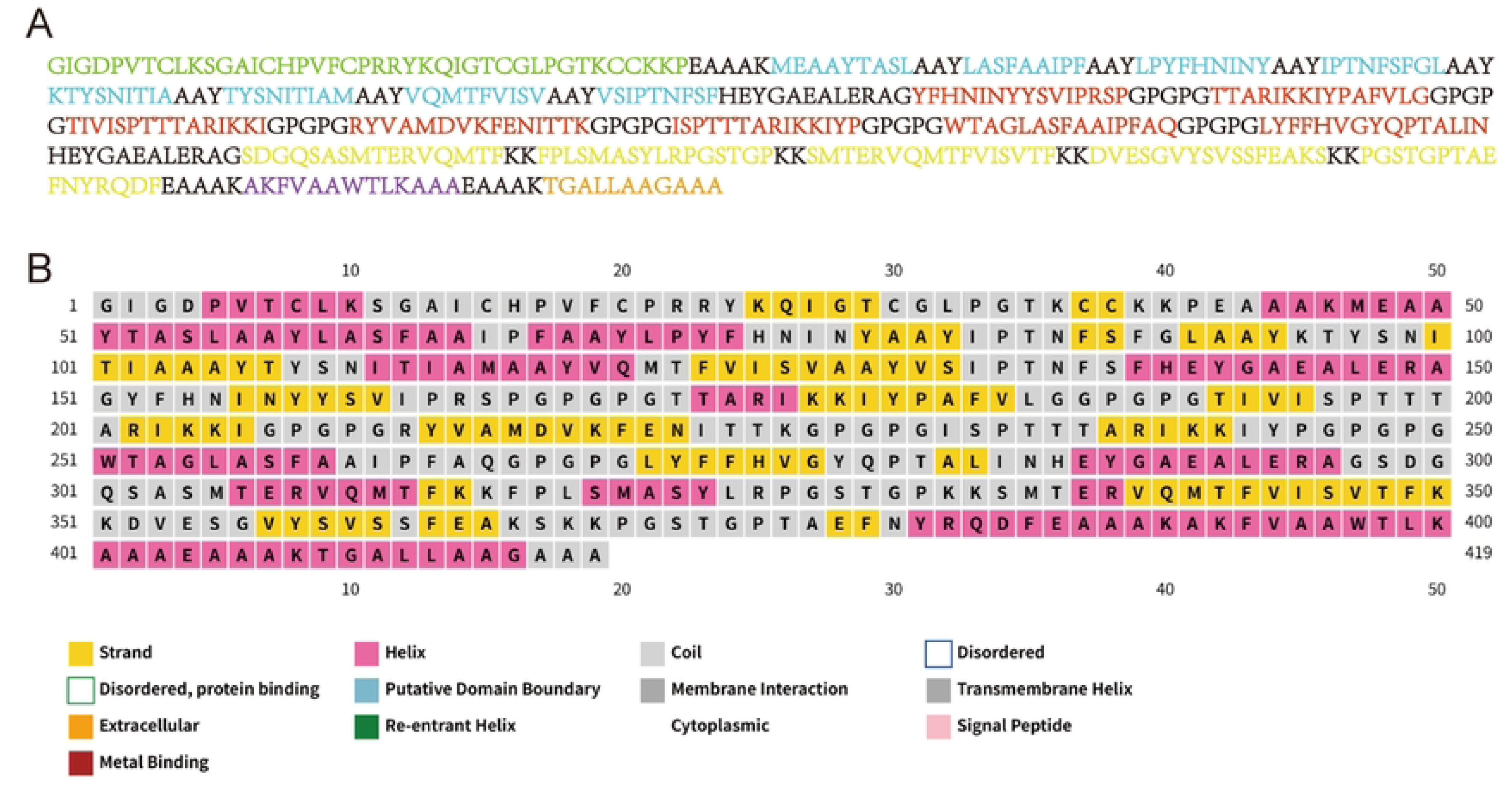
Refined tertiary structure of BtHKU5-CoV-2 vaccine. The model of original (A) and refined (B) structure of the vaccine. (C) The alignment of original construct and refined construct of the vaccine.

**Fig. 3.**
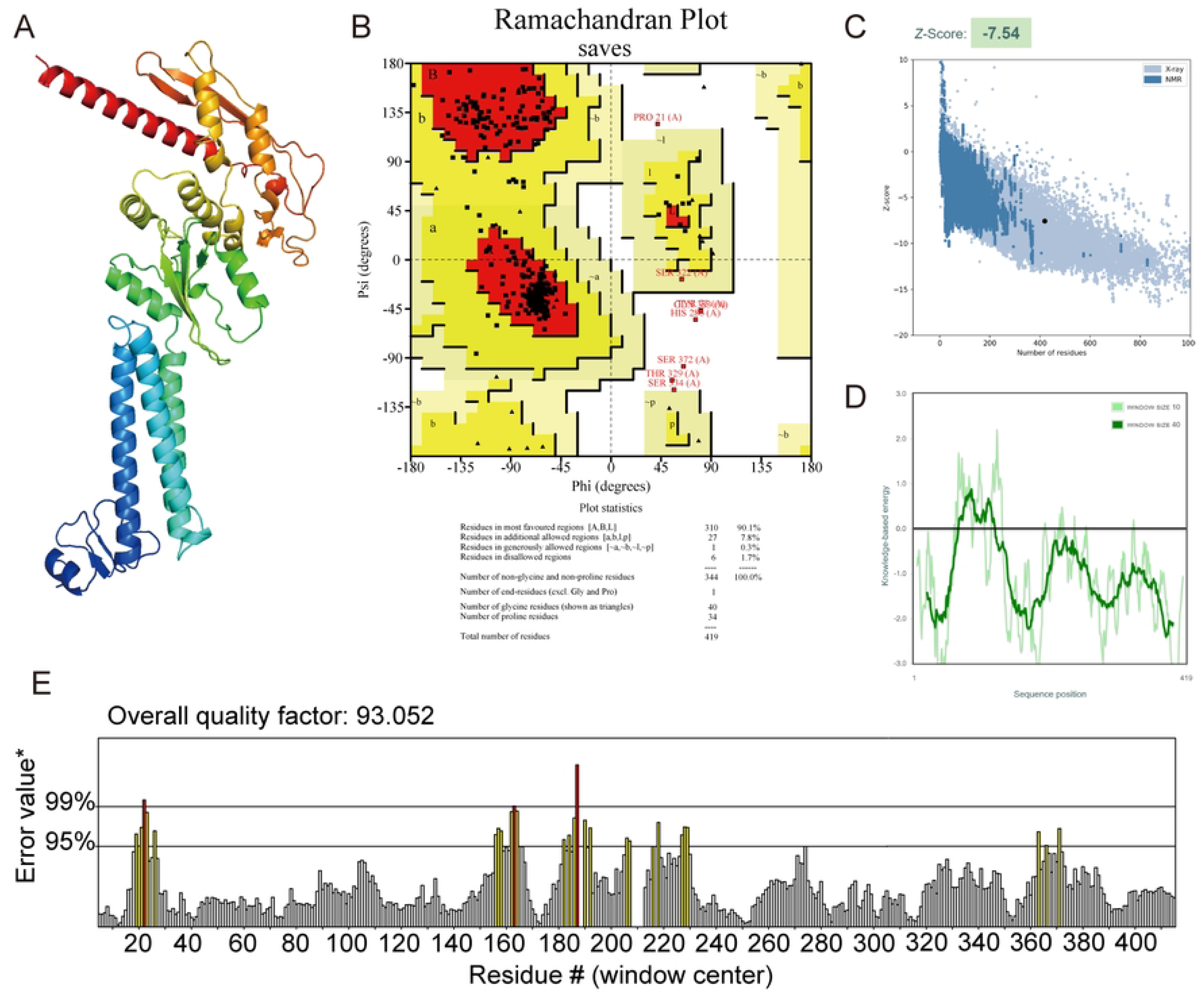
Quality validation for optimized tertiary structure of BtHKU5-CoV-2 vaccine. (A) The refined tertiary structure of the vaccine. (B) The stereochemical quality of the vaccine’s refined tertiary structure was assessed using a Ramachandran plot. In the presented results, red represents the most favoured regions, yellow represents allowed regions and white represents the disallowed regions. (C) Evaluation of the Z-score of vaccine’s optimized tertiary structure was performed by Prosa-Web tools. (D) Energy map of refined structure. (E) In the ERRAT plot, yellow and red represent error values exceeding the 95% and 99% threshold, respectively.

### 4. Prediction of conformational B cell epitopes

To identify conformational B-cell epitopes located in the vaccine’s tertiary structure, the 3D model data were submitted to the ElliPro web server. The results implied that there are four conformational B cell epitopes presented on the surface of the tertiary structure, with scores varied from 0.56 - 0.821 (Table 4). The location of these epitopes in the three-dimensional structure is shown in Fig. 4A-D.

**Fig. 4.**
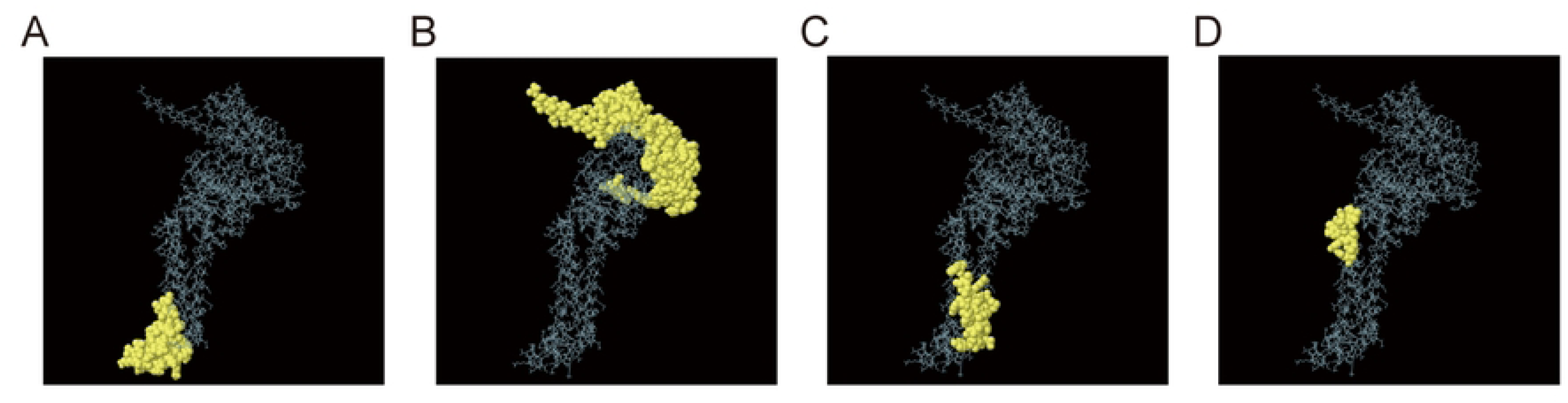
The conformation B epitopes in BtHKU5-CoV-2 vaccine tertiary structure. (A-D) The conformation B epitopes presented in surface model and colored by yellow are located differently in the vaccine’s tertiary structure.

**Table 4.** Conformation B epitope screening results.

| No. | Residues | Number of residues | Score |
| --- | --- | --- | --- |
| 1 | A:G1, A:I2, A:G3, A:D4, A:P5, A:V6, A:T7, A:C8, A:L9, A:K10, A:S11, A:G12, A:A13, A:I14, A:C15, A:H16, A:P17, A:V18, A:F19, A:C20, A:P21, A:R23, A:Y24, A:K25, A:Q26, A:I27, A:G28, A:T29, A:C30, A:G31, A:L32, A:P33, A:G34, A:T35, A:K36, A:C37, A:C38, A:K39, A:K40, A:P41, A:E42, A:A43, A:A44, A:A45, A:K46, A:M47, A:E48, A:A49, A:A50, A:Y51, A:T52, A:A53 | 52 | 0.821 |
| 2 | A:K226, A:G227, A:P228, A:G229, A:P230, A:G231, A:Q265, A:G266, A:P267, A:G268, A:P269, A:A290, A:E291, A:A292, A:L293, A:E294, A:R295, A:A296, A:G297, A:S298, A:D299, A:G300, A:Q301, A:S302, A:A303, A:S304, A:M305, A:T306, A:E307, A:R308, A:Q310, A:M311, A:K314, A:F316, A:P317, A:L318, A:S319, A:A321, A:S322, A:Y323, A:L324, A:R325, A:P326, A:G327, A:S328, A:T329, A:G330, A:P331, A:K332, A:K333, A:S334, A:M335, A:T336, A:E337, A:R338, A:V339, A:Q340, A:M341, A:T342, A:F343, A:V344, A:S346, A:T348, A:K350, A:K351, A:D352, A:V353, A:E354, A:S355, A:G356, A:V357, A:Y358, A:S359, A:V360, A:S361, A:S362, A:F363, A:E364, A:A365, A:K366, A:S367, A:K368, A:K369, A:P370, A:G371, A:S372, A:T373, A:G374, A:P375, A:T376, A:A377, A:E378, A:F379, A:N380, A:Y381, A:R382, A:Q383, A:D384, A:F385, A:W397, A:T398, A:L399, A:K400, A:A401, A:A402, A:A403, A:E404, A:A405, A:A406, A:A407, A:K408, A:T409, A:G410, A:A411, A:L412, A:L413, A:A414, A:A415, A:G416, A:A417, A:A418, A:A419 | 122 | 0.709 |
| 3 | A:R22, A:F89, A:A92, A:A93, A:K95, A:T96, A:Y97, A:S98, A:N99, A:I100, A:T101, A:I102, A:A103, A:A104, A:A105, A:Y106, A:T107, A:Y108, A:S109, A:N110, A:I111, A:T112, A:I113, A:A114, A:M115, A:A116, A:A117, A:Y118, A:M121 | 29 | 0.708 |
| 4 | A:F74, A:H75, A:N76, A:I77, A:N78, A:Y79, A:A81, A:Y82, A:T85, A:N86 | 10 | 0.56 |

### 5. Molecular docking

TLR2 and TLR4 are critical receptors for recognizing viral glycoproteins, thereby activating the immune response [30]. Therefore, binding between the vaccine and these proteins was carried out using ClusPro 2.0. We selected the docking model based on the following criteria: higher cluster rank, lower energy-weighted score, and reasonable binding mode (Table 5). The docking complexes were then refined using the HADDOCK 2.4 tool (Fig.5A and Fig. 6A). The preferred model was based on the HADDOCK score, van der Waals energy, electrostatic energy, and desolvation energy (Table 5). To explore the interaction between the vaccine and the indicated proteins, the PDBsum server was used to analyze the docking complexes (Table 5). Our results revealed that the vaccine formed six hydrogen bonds and one salt bridge with the TLR2 molecule (Fig. 5B), while formed 22 hydrogen bonds and two salt bridges were formed with TLR4 molecule (Fig. 6B). In addition, both the TLR2-vaccine complex and the TLR4-vaccine complex showed very little distortion (Fig. 5C and Fig. 6C). The covariance plot also indicated that both complexes had reasonable dynamic coupling relationships between internal residues (Fig. 5D and Fig. 6D). Overall, these results suggested that the vaccine showed potential stable binding with TLR2 and TLR4.

**Fig. 5.**
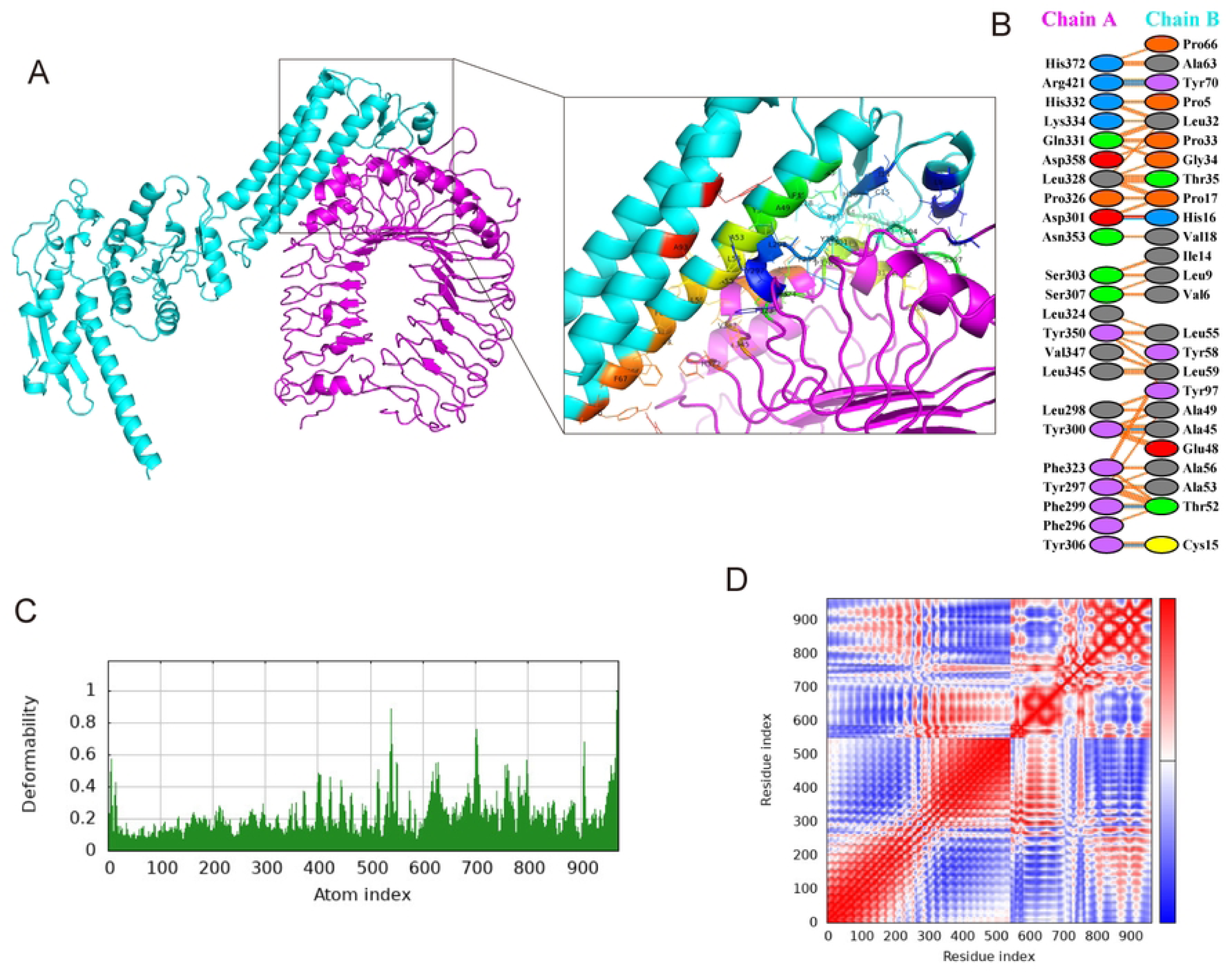
Docking studies of BtHKU5-CoV-2 vaccine-TLR2 molecule complex. (A) The docking complex of HKU5-CoV-2 vaccine and TLR2 molecule. The vaccine is colored in light turquoise, while TLR2 molecule in purple. The boxes show the interaction residues between the vaccine and TLR2 molecules. (B) The connections, including salt bridges, disulphide bonds, hydrogen bonds and non-bonded contacts, identified between amino acids in docking complex, are represented in red, yellow, blue and orange, respectively. (C) Deformability plot of the complex. (D) Covariance plot of the complex. Red, white and blue color indicate correlated motion, non-correlated motion, anti-correlated motion, respectively.

**Fig. 6.**
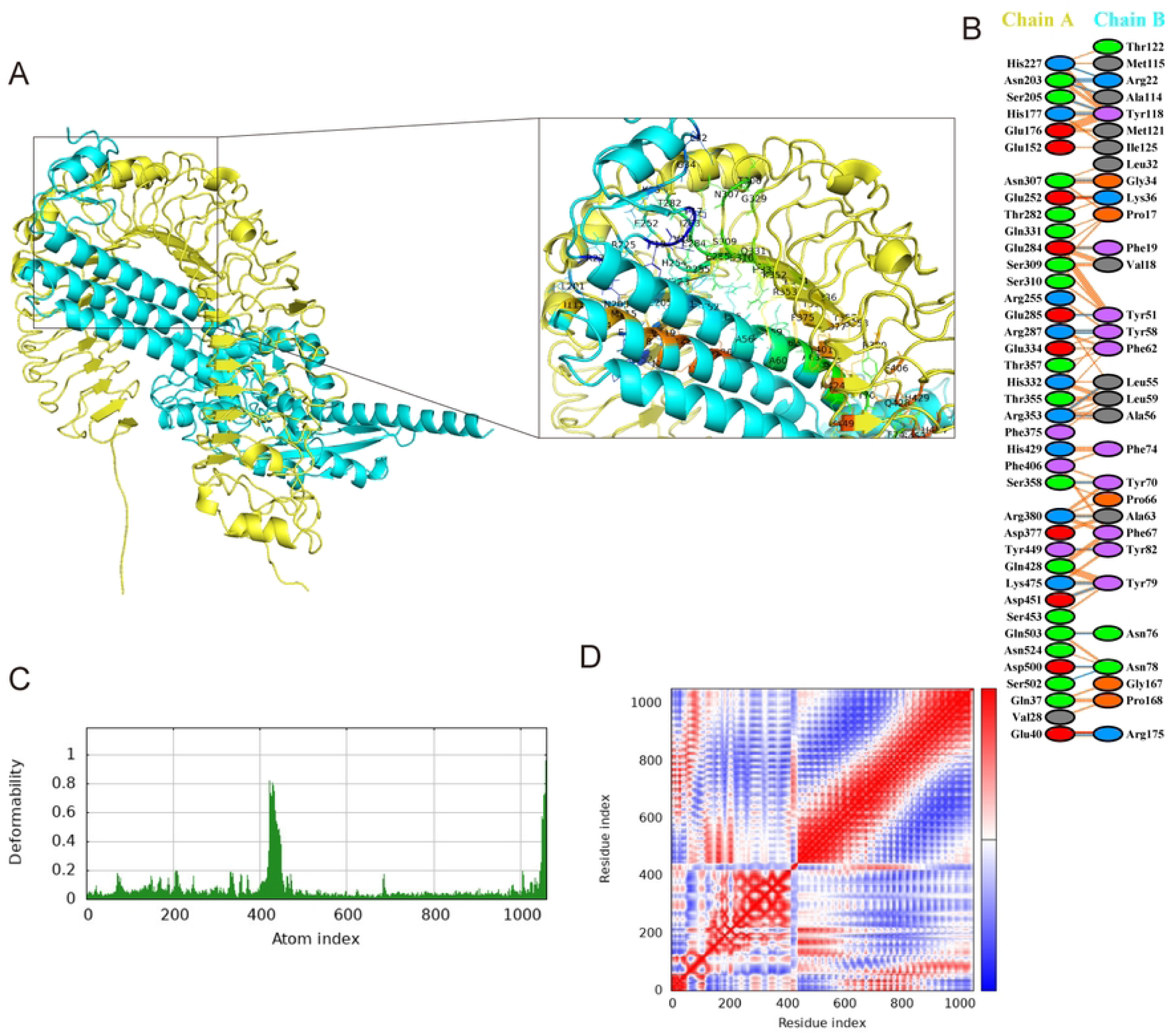
Docking studies of BtHKU5-CoV-2 vaccine-TLR4 molecule complex. (A) The docking complex of HKU5-CoV-2 vaccine and TLR4 molecule. The vaccine is colored in light turquoise, while TLR4 molecule in yellow. The boxes show the interaction residues between the vaccine and TLR4 molecules. (B) The connections, including salt bridges, disulphide bonds, hydrogen bonds and non-bonded contacts, identified between amino acids in docking complex, are represented in red, yellow, blue and orange, respectively. (C) Deformability plot of the complex. (D) Covariance plot of the complex. Red, white and blue color indicate correlated motion, non-correlated motion, anti-correlated motion, respectively.

**Table 5.**
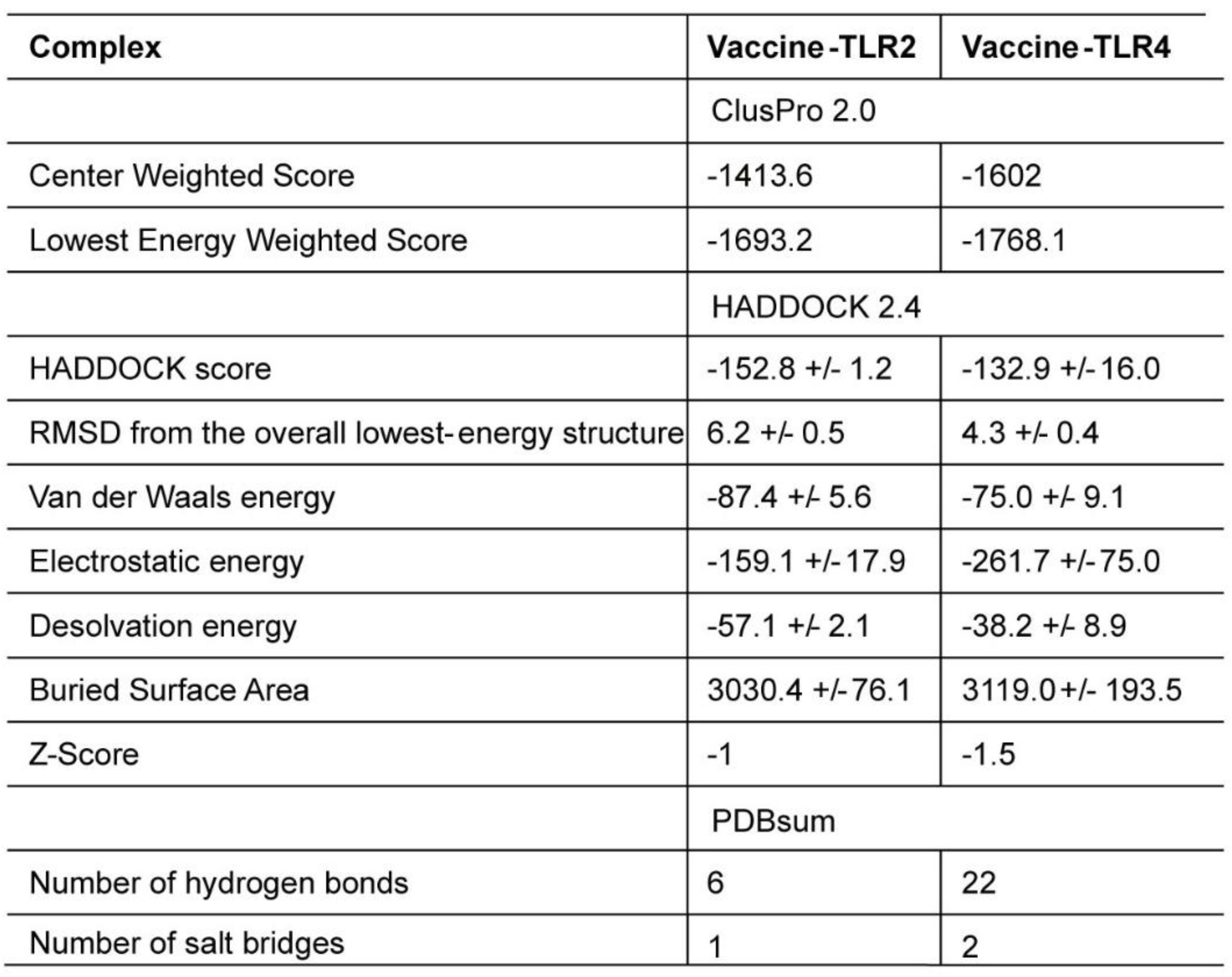
Results of molecular docking.

| Complex | Vaccine-TLR2 | Vaccine-TLR4 |
| --- | --- | --- |
|  | ClusPro 2.0 |  |
| Center Weighted Score | -1413.6 | -1602 |
| Lowest Energy Weighted Score | -1693.2 | -1768.1 |
|  | HADDOCK 2.4 |  |
| HADDOCK score | -152.8 +/- 1.2 | -132.9 +/- 16.0 |
| RMSD from the overall lowest-energy structure | 6.2 +/- 0.5 | 4.3 +/- 0.4 |
| Van der Waals energy | -87.4 +/- 5.6 | -75.0 +/- 9.1 |
| Electrostatic energy | -159.1 +/- 17.9 | -261.7 +/- 75.0 |
| Desolvation energy | -57.1 +/- 2.1 | -38.2 +/- 8.9 |
| Buried Surface Area | 3030.4 +/- 76.1 | 3119.0 +/- 193.5 |
| Z-Score | -1 | -1.5 |
|  | PDBsum |  |
| Number of hydrogen bonds | 6 | 22 |
| Number of salt bridges | 1 | 2 |

### 6. Population coverage calculation

To estimate the percentage of individuals the vaccine candidate could provide protection across the world, the vaccine’s population coverage was calculated using the IEDB database. As shown in Fig. 7, approximately 97.49% of the global population benefits from vaccination. In particular, the vaccine achieved coverage rates of 99.99% in the United States, 94.65% in China, 99.81% in France, and 99.78% in Russia.

**Fig. 7.**
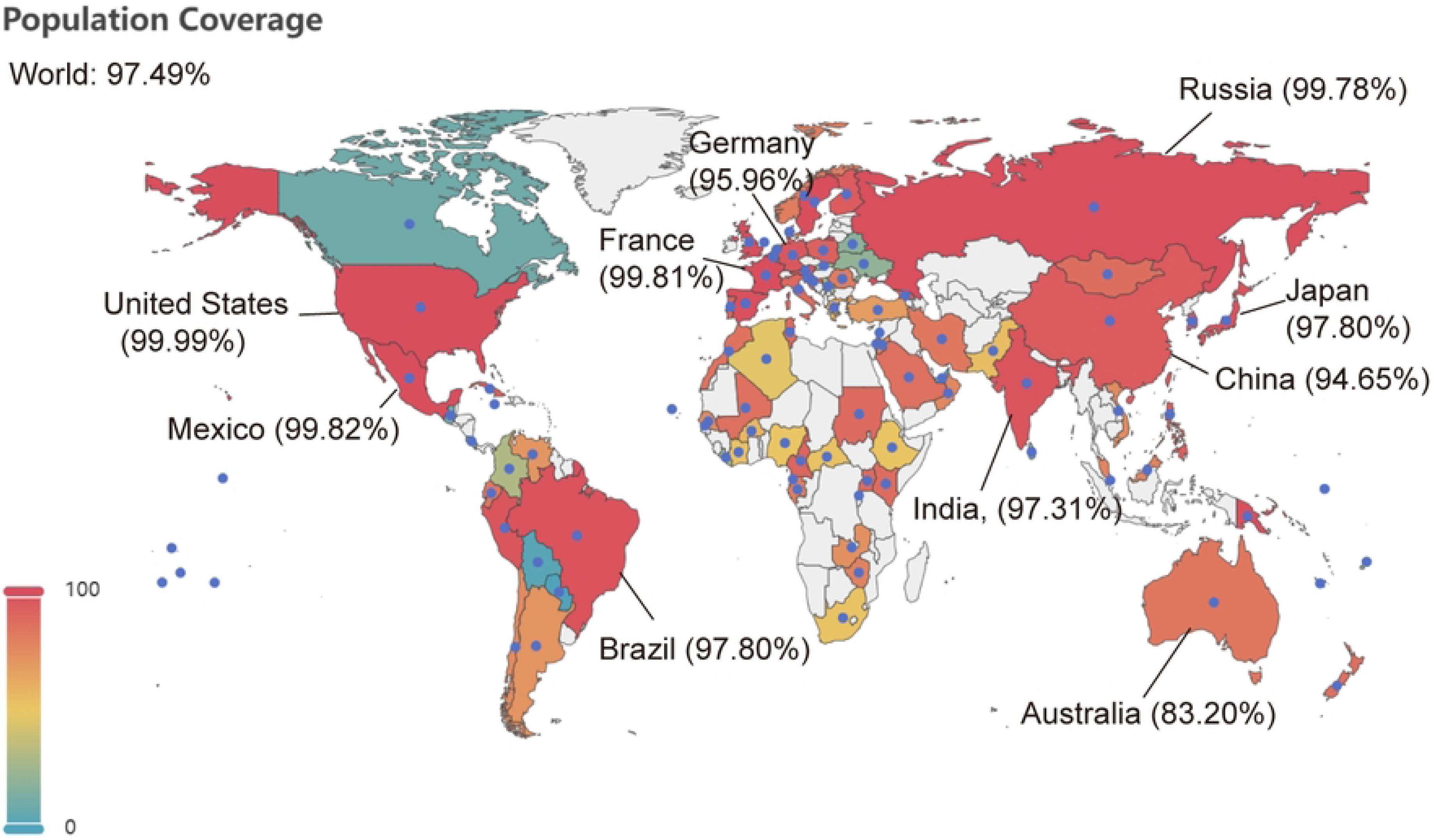
Population coverage of BtHKU5-CoV-2 vaccine across the world. The countries and regions that have available data are labeled with blue dots.

### 7. In silico immunization simulation

To better map the dynamic process of immune response within the host organism post vaccination, the C-IMMSIM server was used to assess the effect of the mRNA vaccine. This server depicts the immunogenic profile after three vaccine injections for almost one year. As shown in Fig. 8A, the antibody levels were significantly increased following the third injection, with the level of IgM plus IgG peaking at approximately 1.6 × 10^5^. Although IFNg and IL-2 were released once the vaccine was administered, they were quickly cleared from the plasma (Fig. 8B). During the simulation period, the populations of B cells, plasma cells, and helper T cells initially showed marked expansion following injection, and then gradually decreased (Fig. 8C-E). A large number of resting cytotoxic T cells (TC) were activated and transformed into active cytotoxic T cells when the multi-epitopes stimulated the host immune system, although a gradual decline in active TC cells was observed after the third vaccine introduction (Fig. 8F). The total populations of TC cells, natural killer (NK) cells, and dendritic cells (DCs) consistently increased following administration of the first dose of the vaccine, and eventually remained in equilibrium in a dynamic range (Fig. 8G-I). These results indicate that the mRNA vaccine used in the present study can elicit a robust adaptive immune response in the host.

**Fig. 8.**
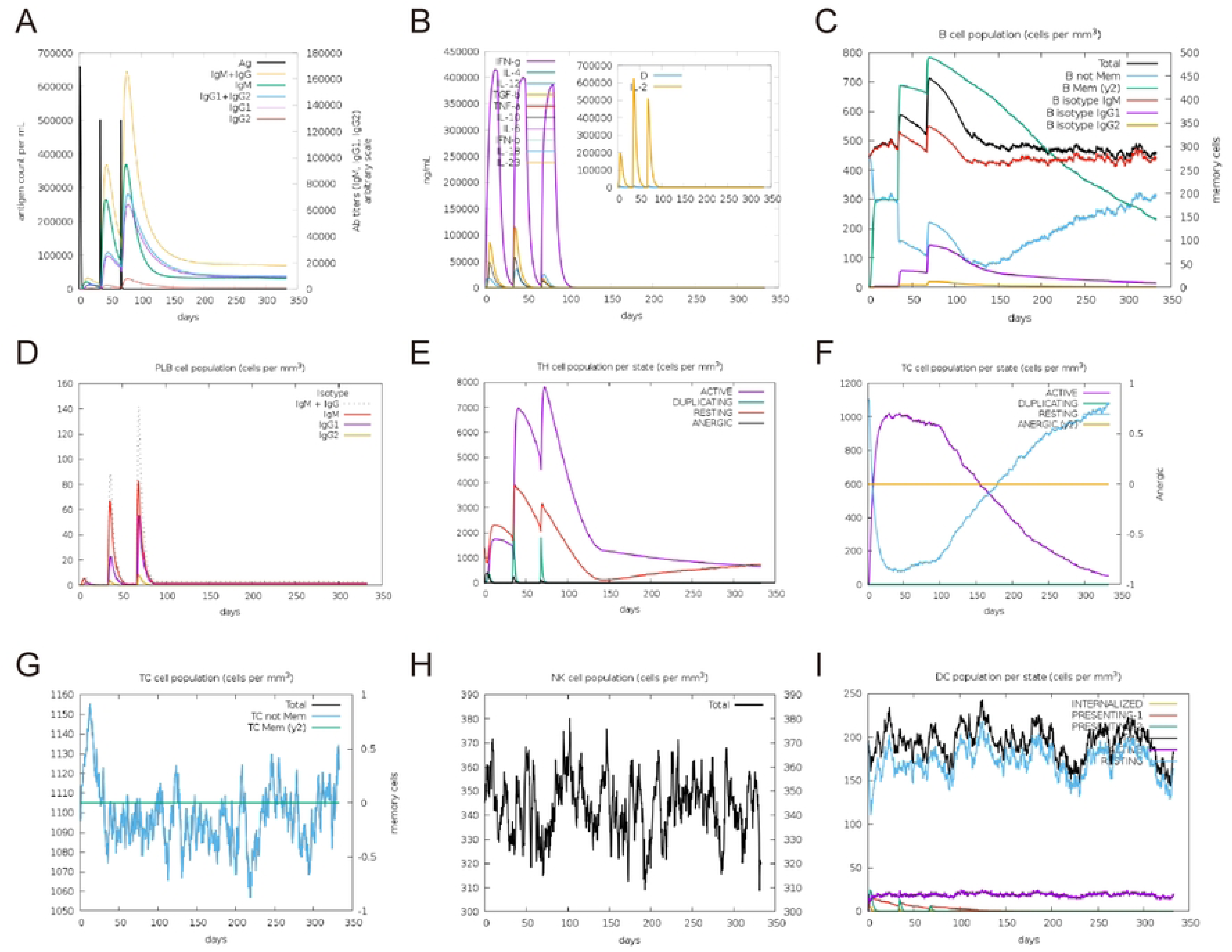
Immunological simulation analysis after administration of three doses of BtHKU5-CoV-2 vaccine. Various types of immunoglobulin (A) or cytokines (B) were produced following vaccine injection. The levels of B cells (C), plasma cells (D), helper T (TH) cells (E), TC cells (F and G), NK cells (H) and DCs (I) at different states were depicted post vaccination.

### 8. mRNA vaccine construction

After attaching the tPA and MITD sequences to the N-terminus and C-terminus of the vaccine, respectively, the full-length amino acid sequence of the final mRNA vaccine was submitted to the GenSmart Codon Optimization tool for reverse translation and code optimization. The improved cDNA fragment showed a Codon Adaptation Index (CAI) value of 0.91 and a 61% GC content, indicating that the vaccine exhibits high expression in humans (S2 Fig). Upon merging the 5’-UTR, 3’-UTR, and poly(A) tail, the second structure of the full-length RNA sequence was analyzed using an RNA fold server (S3 Fig). The free energy of the thermodynamic ensemble was predicted to be -678.61 kcal/mol, suggesting a stable RNA structure.

TLR3, TLR7, and TLR8 are critical sensors of exogenous nucleic acids and are located in endosomes [34]. We first predicted the binding between the vaccine’s full-length mRNA and these TLRs molecules using the RPISeq tool based on RF and SVM methods. All binding scores exceeded 0.5, suggesting that the vaccine mRNA may interact with TLR3, TLR7, and TLR8 (S1_Table). We then ran a molecular docking analysis for TLR3, TLR7, and TLR8 using the HDOCK server. The docking models are shown in Fig. 9A-C, and the binding and confidence scores are presented in Table. S1B. All binding scores were above 200 with high confidence scores, indicating that the vaccine mRNA might be able to interact with TLR3, TLR7, and TLR8 molecules.

**Fig. 9.**
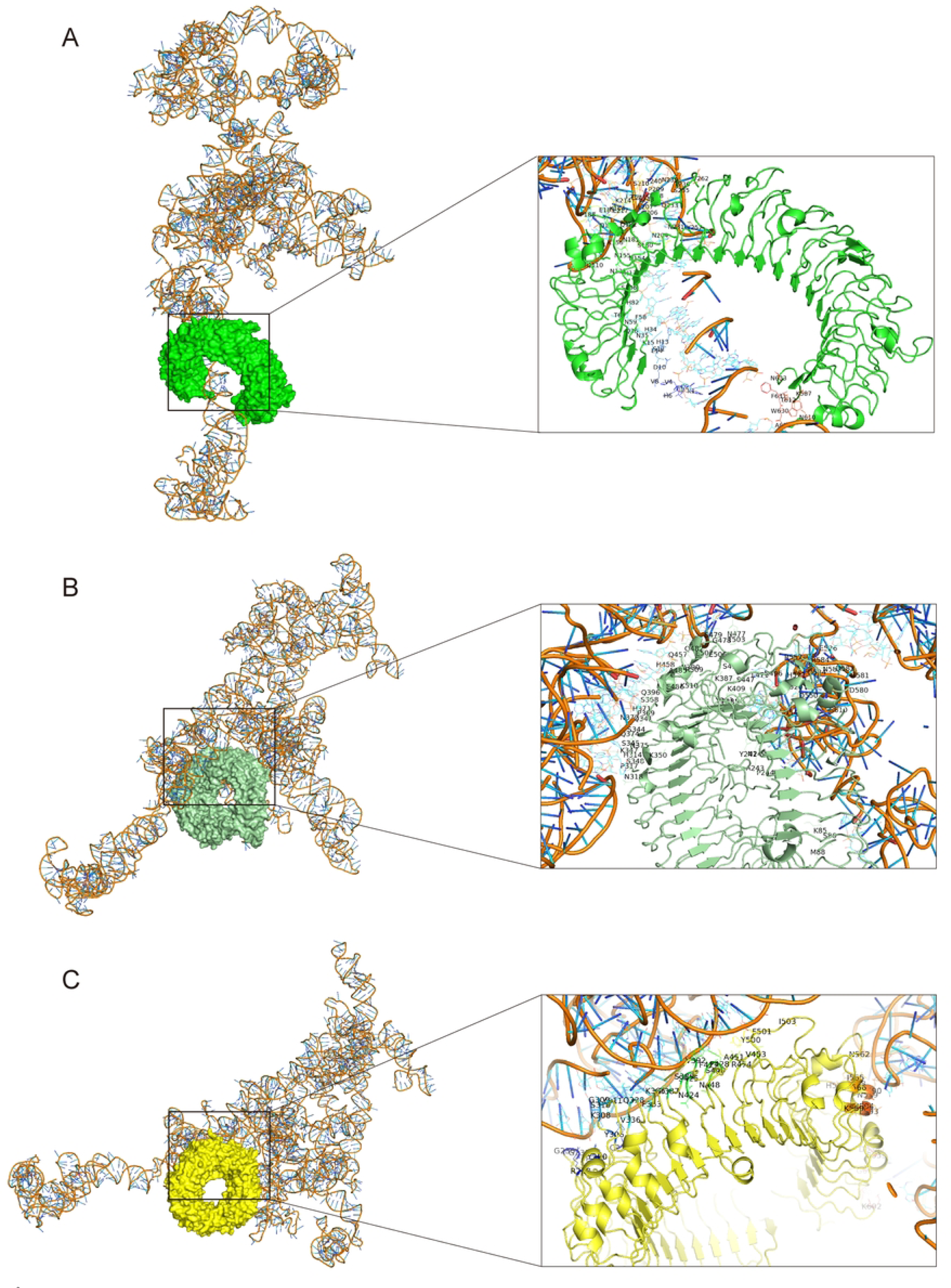
Docking analysis of BtHKU5-CoV-2 vaccine (mRNA)-TLR molecules complexes. The docking models of the vaccine mRNA with TLR3 molecule (A), TLR7 molecule (B) or TLR8 molecule (C) was showed. In these complexes, the mRNA is presented as complicated strand, while TLR3 is presented as red-colored half ring, TLR7 as light-colored ring and TLR8 as yellow-colored ring. The boxes show the interaction residues between the mRNA and TLR molecules.

Finally, the full-length DNA of the mRNA vaccine was introduced into the pET-28a (+) vector via homologous recombination (S4 Fig). The plasmid was presumed to act as a template for in vitro transcription, and the synthesized mRNA was encapsulated by lipid nanoparticles and subjected to vaccination.

## Discussion

Since the first discovery of BtHKU5-CoV-2 in March 2025, widespread concerns have been raised [5–7]. BtHKU5-CoV-2 was found to infect human cell lines by utilizing the human ACE2 receptor, similar to SARS-CoV-1 and SARS-CoV-2, which led to severe acute respiratory syndrome (SARS) and the COVID-19 pandemic [1, 35, 36]. Moreover, it adapts to ACE2 orthologs of many mammalian and avian species, indicating that it might have a wide host range [1]. Although no case of human infection has been reported, BtHKU5-CoV-2 has the potential to spill over to humans and cause a pandemic outburst. Although the team of Prof. Zhengli Shi demonstrated that some protease inhibitors (such as E64d), endosomal acidification inhibitors (such as bafilomycin A1), one fusion inhibitor EK1C4, and several small molecule inhibitors (including nirmatrelvir and remdesivir) suppressed the entry and replication of BtHKU5-CoV-2 in vitro [1], it is necessary to develop prophylactic and therapeutic vaccines against BtHKU5-CoV-2. Engagement of host receptor recognition and viral and cell membrane fusion is mediated by spike glycoprotein on the surface of coronaviruses [37]. Therefore, the spike glycoprotein is a critical target for the development of prophylactic vaccines. During response to outbreaks or rapidly evolving pathogens, nucleic acid vaccines present more advantages than traditional vaccines, including inactivated vaccines and subunit vaccines [38]. The success of combatting COVID-19 has accelerated the development of mRNA vaccines.

In the present study, we designed a multi-epitope mRNA vaccine to protect against BtHKU5-CoV-2 infection. The epitope sequences were obtained from the conserved regions of the spike glycoprotein of BtHKU5-CoV-2, ensuring that the multi-epitope vaccine could cover all six BtHKU5-CoV-2 strains. Eight CTL epitopes, seven HTL epitopes, and five LBL epitopes with strong antigenicity and immunogenicity were selected for vaccine construction. These epitopes were expected to trigger antibody-mediated immunity and T-cell immune responses, but did not cause autoimmune diseases. This is because epitopes show no homology with proteomes of human and commensal bacteria in the human intestine. In accordance with previous studies, linkers AAY, GPGPG, KK, and EAAAK were used to connect the indicated epitopes [39, 40]. Incorporating linkers was expected to facilitate accurate cleavage and proper folding of the multi-epitope vaccine. Beta-defensin II acts as a ligand for TLRs receptors, thereby activating the immune response [41, 42]. Thus, β-defensin II was attached to the N-terminus of the vaccine construct and acted as an adjuvant. Furthermore, a cell entry peptide, TAT sequence, was added to the C-terminus of the vaccine construct, and it was proposed to target vaccines across the cellular membrane [43].

To further determine whether the protein encoded by the mRNA vaccine posed a proper folding conformation, we predicted the tertiary structure using the Robetta server. The rationality of the refined tertiary structure was assessed using a Ramachandran plot, and the results indicated the good quality of the spatial structures. A comprehensive understanding of the interaction between vaccines and immune receptors, especially TLR2 and TLR4, is crucial for vaccine development. Molecular docking analysis showed that the β-defensin II domain of the vaccine exhibited strong docking with TLR2 and TLR4. In addition, hydrogen bond and salt bridge analyses indicated that the refined vaccine tightly interacted with TLR2 and TLR4, suggesting the potential of the vaccine candidates to induce an immune response.

Previous reports on SARS-CoV-2 have demonstrated that SARS-CoV-2 RNA is recognized by TLR3, TLR7, and TLR8, resulting in the activation of innate immune signaling and production of cytokines and interferons [44, 45]. The RNA-protein interaction prediction scores obtained in this study suggested that the mRNA of the designed vaccine showed high binding affinity with TLR3, TLR7, and TLR8 molecules. Therefore, the vaccine mRNA may assist in triggering the host immune response.

Finally, the immune response induced by the developed vaccine was predicted using C-IMMSIM server. On the one hand, high levels of antibodies, which play critical role in preventing BtHKU5-CoV-2 infection, were observed after the vaccine injection. On the other hand, vaccination significantly increased the number of helper T cells and active cytotoxic T cells in the peripheral blood, indicating strong cellular immunity. Furthermore, the vaccine had the potency to induce long-lasting immunity because the antibodies, including IgG and IgM, tended to remain stable for a long period of time, while memory B cells and memory T cells were generated after administration of three doses of vaccines. Collectively, the results of our study strongly support that the designed mRNA vaccine has the potential to induce the desired humoral immunity and cellular immunity and will be a preferred candidate for preventing future outbreaks of BtHKU5-CoV-2.

## Author contributions

Ningze Zheng: designed the study, performed data processing and wrote the manuscript draft.

Yingqi Xu: performed data processing and wrote the manuscript draft.

All authors have read and approved the final manuscript. All authors agree to be accountable for all aspects of the work.

## Data availability statement

The authors confirm that the raw data supporting the findings of this study are openly available in Science Data Bank at https://doi.org/10.57760/sciencedb.28908.

## Supporting information

S1 Fig. Vaccine’s solubility (QuerySol) was predicted through Protein-Sol tool.

PopAvrSol represents the average solubility of the proteins in experimental dataset.

S2 Fig. Codon adaptive index (CAI) analysis of the coding regions of BtHKU5-CoV-2 vaccine.

The CAI and GC content of the vaccine are 0.91 and 61%, respectively.

S3 Fig. Secondary structure of BtHKU5-CoV-2 mRNA.

(A) The MFE secondary structure for minimum free energy prediction. (B) The centroid secondary structure for thermodynamic ensemble prediction.

S4 Fig. Schematic structure of BtHKU5-CoV-2 vaccine plasmid.

The inserted sequence indicated the full-length DNA of HKU5-CoV-2 mRNA vaccine.

